# Towards Quantitative *In Vitro* Modelling of Focused Ultrasound-mediated Blood-Brain Barrier Opening using an Ultrasound-transparent Organ-on-Chip Device

**DOI:** 10.64898/2026.09.29.755358

**Authors:** Nabhan M. Fakrudin, Nathan Han, Laith Kabbani, Brandon Lan, Suzan Aldekr, Baiju Thomas, Matthew W. Barrett, Rayna J. Gonzales, Frederic Zenhausern, Jian Gu

## Abstract

Gas-bubble-enhanced focused ultrasound (FUS^GB^) is advancing clinically across a myriad of applications, including targeted drug delivery through transient blood-brain barrier opening (BBBO). Its broader translation requires a better understanding of the bioeffects that enable reproducible enhancement of barrier permeability while avoiding vascular injury, underscoring the need to clarify the relationship between acoustic exposure, bubble activity, and the biological response central to treatment optimization. These interactions are difficult to isolate in vivo, and acoustic reflections, geometry, and field distortion in conventional cultureware hinder quantitative in vitro studies. Herein, we present a theoretically and experimentally characterized ultrasound-transparent organ-on-chip device with >99.9% US transparency that enables quantitative in vitro modeling of the FUS^GB^ procedure and study of BBBO. Furthermore, submicron bubbles (sMB, ~800nm) were fabricated and characterized. Acoustic dose-dependent subharmonic and broadband cavitation dose curves (SCD and BCD), as well as temporal responses, were studied. A 3-day accelerated Caco-2 surrogate barrier with a high TEER (~1200 O·cm2) was used to demonstrate US dose-dependent targeted barrier opening, visualized using an in vitro Evans Blue assay. Together, this study and its findings establish a foundation for quantitative in vitro modeling of FUS BBBO, as well as other US interactions in microphysiological systems, supporting systematic evaluation of procedural regimens, large-parameter-space optimization, acoustic safety window, and exposure conditions for brain drug delivery, and additionally, enabling mechanistic insight into FUS^GB^ BBBO.

## Introduction

Ultrasound (US) plays a pivotal role in medicine, not only in imaging^1^ but also in a growing range of therapeutic applications, such as cancer^2–5^, cardiovascular disease treatment^6^, gene therapy & transfection^7,8^, drug delivery^9^, ischemic stroke rehabilitation^10^, and others. Given its high degree of non-invasiveness for the aforementioned applications and more, it is a unique next-generation technique for surgical and drug-delivery procedures. Developing safe and effective US therapies and treatments depends on understanding their bioeffects on cells and tissues under well-controlled conditions, as these effects can be complex.^11–13^ Currently, animal models are widely used for therapeutic US preclinical studies; however, animal testing can be limited by throughput, cost, ethical considerations, attrition during translation due to species differences, and the predictability of clinical trial outcomes.^14–17^

Recently, organ-on-chips (OoCs), also known as microphysiological systems (MPS), have shown promise as alternatives to animal testing for certain use cases.^14,15,18,19^ OoCs are purpose-engineered in vitro cell systems that replicate human tissue and organ structures and their physiology to study physiology, etiology, and disease treatment.^20,21^ Nonetheless, investigating the bioeffects of US on and in OoC systems (or in vitro systems in general) presents considerable challenges due to multiple factors, including reflective interfaces, inconsistent fluid volumes in the acoustic path^22^, and material properties that can lead to significant acoustic impedance mismatches between commonly used in vitro cell culture vessels and the aqueous environment that supports cellular life.^23,24^ To that end, Hensel *et al*. have experimentally shown that reflective interfaces can shift local pressure maxima, increasing them by up to a factor of 5, and that minor variations as low as 2.56% in fluid volume along the acoustic path can change the acoustic dose experienced by the cells by up to a factor of two.^22^ Furthermore, Leskinen *et al*. experimentally validate that US energy uncertainties can reach as high as 700%, attributable to impedance mismatch^25^, and Gupta *et al*. speculate, based on observed vibration displacement, an acoustic exposure variance for the cells between 110-120% and 58-80% in 6-well plates and Petri dishes, respectively.^23^ As a result, large variation and poor reproducibility are often encountered for in vitro US experiments^26^, which also hinders the development of OoC models for therapeutic US applications. For example, Beekers *et al*. reported using high-speed microscopy and microbubble response to characterize the acoustic field inside an OoC device. Values up to 10 dB away from free-space pressure and interquartile pressure variation up to 7 dB have been reported^27^, and other studies have similarly reported in situ pressure variance of up to −8.7 to −1.1dB in a thickness-dependent manner and an attenuation of up to 65.5% of the incident pressure.^28^ Several in vitro studies have since reported promising bioeffects and observations^29–39^; however, these models are limited by transwell-based architecture with through-plate insonication, or, in models using PDMS or other materials, impedance mismatches and reflective boundaries persist.^34,37,38,40,41^

One known approach to study US interaction in Transwell inserts and minimize attenuation of the incident wave is the direct immersion of the transducer into the insert; however, this method suffers from reflective boundaries at the plate-liquid interface and in the surrounding plastic, leading to standing waves that can influence bioeffects and, by some accounts investigating sonoporation regimes, make them poorly reproducible in vivo.^42,43^ A newer method overcomes this by inverting the Transwell directly above the transducer and incorporating an acoustic matching boundary layer (Sorbothane® foam) at the air-liquid interface beyond the insert to minimize reflections.^30,44–46^ This approach is particularly useful for examining the bioeffects of larger gas bubbles (>3-6µm), which can rise at velocities up to 20 µm/s (<1 min/mm rise time) to remain in contact with the cell layer during insonication; however, it is not pragmatic for high-throughput studies assessing US exposure safety and recovery for patient-derived models, given the obvious risk of handling-induced variation in the cellular barrier^46^, risk of biological contamination making results unreliable, and effort needed.

A method for accurately delivering US energy to in vitro cell cultures uses a thin plastic film-like PET, polystyrene, or Parafilm™-to create a US-transparent (UST) window that improves transmission and reduces reflection and standing-wave formation.^47^ According to the theory of acoustic wave propagation through layered media, the thin film is expected to become UST as its thickness approaches zero.^48^ However, to our knowledge, this has not yet been applied to OoC platforms.

Herein we describe a UST-OoC device made with polyethylene terephthalate (PET) thin films and a porous PET membrane within a bilayer polycarbonate (PC) channel architecture, commonly used for apo-basolateral barrier research. The transmission and reflection properties of the PET thin films were measured at different angles of incidence (AOI) and compared with theoretical models. An OoC device featuring the UST window was demonstrated, and acoustic field simulations were performed to evaluate field uniformity and US-dose exposure within the device. We then produced submicron bubbles (sMBs) (between 500nm and 1µm) and evaluated their dose-dependent acoustic behavior using sub-harmonic cavitation dose (SCD) along with broadband cavitation dose (BCD). sMB-assisted focused ultrasound (FUS^sMB^) was then used to disrupt an accelerated Caco-2 cellular barrier within the UST OoC device. Dose-dependent, targeted barrier opening with spatial resolution was visualized using an in vitro Evans Blue assay, demonstrating the UST-OoC device’s effectiveness for studying dose-dependent ultrasound effects in vitro.

## Results & Discussion

### Theoretical analysis and acoustic characterization of PET thin films in water

Acoustic transmission and reflection coefficients were modeled for a uniformly propagating medium 1 with an acoustic impedance of Z_1_, incident on a thin film of medium 2, with thickness (d) and acoustic impedance Z_2_ (**Fig. 1a**). The AOI and refraction are denoted by θ_1_ and θ_2_, respectively. According to the acoustic wave theory described by L. M. Brekhovskikh^48^, the transmission coefficient *t* and reflection coefficient *r* of the incident wave onto the stack of thin films are:

**Figure 1:**
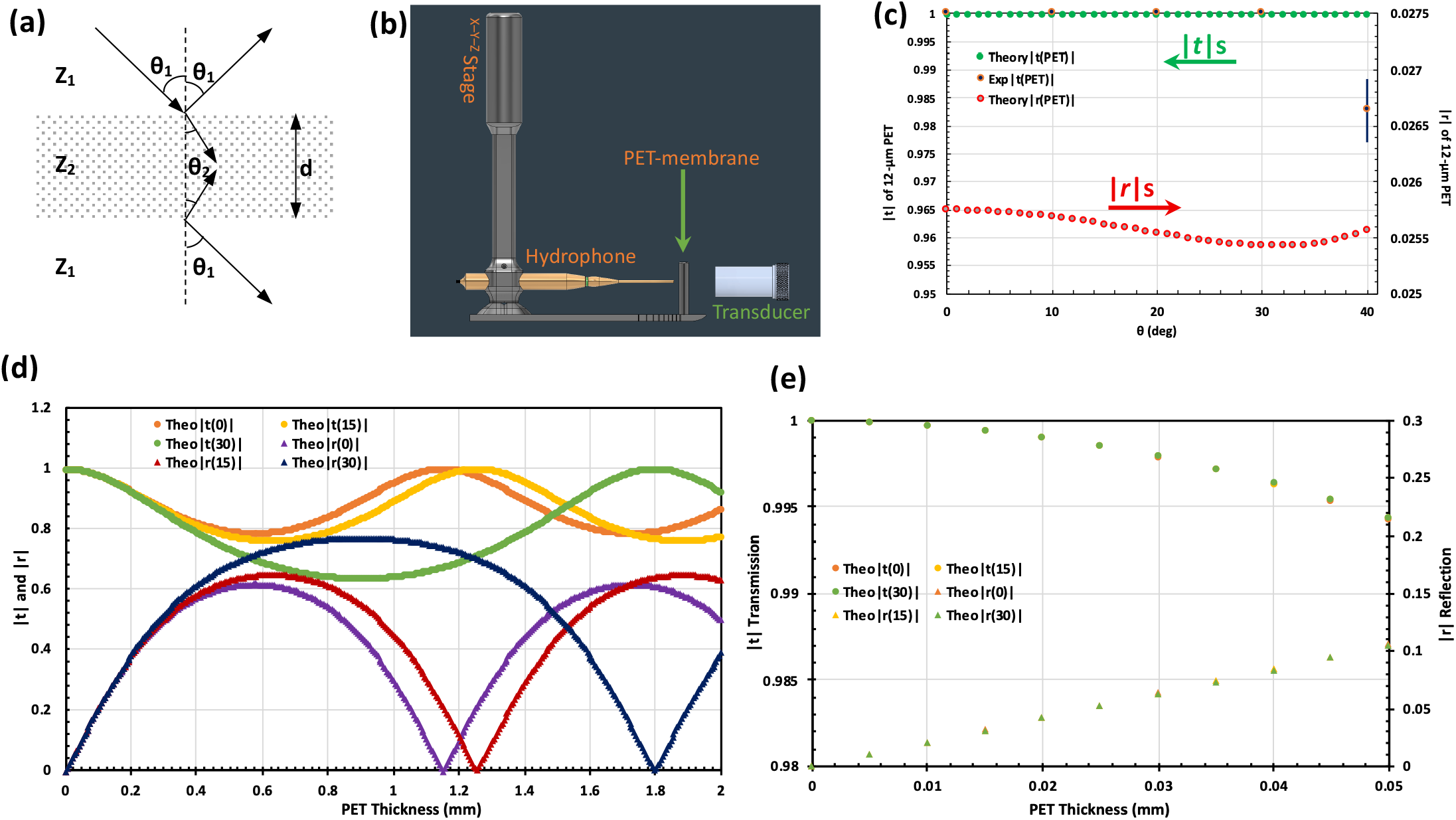
**a)** Schematic of an incident acoustic wave on a thin film; **b)** Rendering of the custom setup used for measuring the |*t*| and |*r*| coefficients; **c)** Theoretical |*t*| and |*r*| with experimental |*t*| measurements for the PET thin film used at different AOI(∅); **d)** (left) Solution to the theoretical calculation of |*t*| and |*r*| for a PET thin film of different thicknesses at different angles of incidence; (right) a zoomed-in view for a PET thickness of 0-50µm.

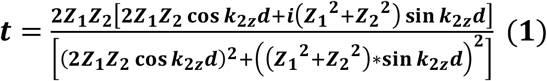

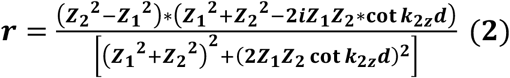

where the wave vector 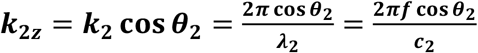, and the impedances 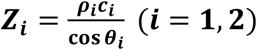, *f* is the wave frequency *λ*_*i*_, *c*_*i*_ and *ρ*_*i*_ are the wavelengths, sound speeds, and densities of the respective media. From Eq. (1–2), the amplitudes of *t* and *r* can be further derived as:

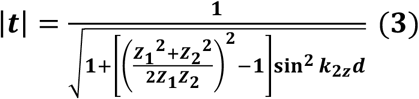

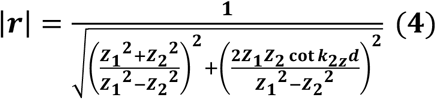

Eq. (3–4) shows that when ***d*** → **0**, | ***t*** | → 1 ***and*** | ***t*** | → **0**, transmission improves, and reflection decreases, minimizing standing-wave formation.

To assess how well the theoretical calculation applies to a PET thin film in water for fabricating UST-OoC devices, we mounted a 12-µm PET thin film on a polycarbonate (PC) frame with a 13-mm center hole to form a PET thin-film window. A 1 MHz water-immersion FUS transducer was used to generate a pulsed US signal directed at the center of the PET thin-film window at angles of incidence of 0°, 15,° and 30,° and the transmission coefficient was measured with a 1-mm needle hydrophone. For reflection coefficient measurements, we used a 45°AOI due to geometric limitations. **Fig. 1b** shows the setup rendering, where the transducer, PET thin film, and needle hydrophone were submerged in a tank filled with degassed deionized (DI) water. **Fig. S1 (Supplemental Materials)** shows images of the actual setup for transmission and reflection measurements, and the electrical driving setup used.

**Table 1** lists the acoustic properties of water and PET for the theoretical calculation. **Fig. 1d-e** shows the theoretical solutions for how the amplitudes of *t* and *r* change with PET thin-film thickness. The top curves show that the theoretical |*t*| value varies periodically with PET film thickness, with a wave period of 1,150 µm at normal incidence for a 1 MHz wave in water and a minimum |*t*| of 78.6% at half the period (575 µm). This behavior aligns with the half-wave layer effect, in which the membrane does not affect the incident wave’s transmission because out-of-phase reflections cancel each other out, bringing |*t*| close to unity.^48^ The period increases to 1,255 µm and 1,795 µm for angles of incidence of 15 and 30 degrees, with minimum |*t*| of 76.1% and 63.7%, respectively. The shift and broadening of the observed resonance can be attributed to the increased effective path length. Indeed, when PET film thickness goes to zero, |*t*| approaches unity for all AOIs, i.e., full transmission.

**Table 1:** Acoustic properties of water and PET used for theoretical calculation and simulation at 1MHz.

| Materials | Water | PET |
| --- | --- | --- |
| Density $\rho$ ($10^3 \text{ kg/m}^3$) | 0.997<sup>49</sup> | 1.335<sup>50</sup> |
| Longitudinal wave sound speed $c$ (km/s) | 1497<sup>49</sup> | 2300<sup>51</sup> |
| Acoustic impedance $Z=\rho c$ (MRy) | 1.493 | 3.071 |

The theoretical solutions were experimentally tested for a 12 µm-thick PET membrane across 0-40°AOI (**Fig. 1c**). Experimentally, the |*t*| results show minimal deviation from theoretical predictions and near-unity transmission up to 30°AOI; however, a slight drop of 1.68% was observed at an AOI of 40°, possibly due to the reduced UST window size at increased AOI.

### Fabrication of an OoC device with a UST window

A key objective in developing the UST OoC was to characterize and demonstrate US transparency with minimal scattering, reflection, and absorption to enable accurate US power delivery. Once we confirmed the experimental feasibility of an ultrasound-transparent window made of PET thin film (previous section), with high transmission (|*t*|>99.96%) and low reflection (|*r*|<2.57%) for a ~12 µm PET film, we next evaluated the incorporation of the windows within a bilayer apo-basolateral sandwich architecture device commonly used for barrier studies^52^. The device is built around PET thin films as the capping layers of the apical and basolateral channels, with a central 12 µm-thick porous (3 µm pore dia.) hydrophilic membrane that allows culture of a cellular barrier (**Fig. 2a, b**). This architecture enables PET-media-water coupling to minimize gross media-impedance mismatch and reflections while maintaining an isolated, sterile environment within the device, thereby providing a contamination-free setting for recovery and long-term studies. Another key design benefit is that this architecture can scale to large-scale, high-throughput studies, enabling evaluation of personalized safety regimens. The methods section of the manuscript provides more detail on the fabrication.

**Figure 2:**
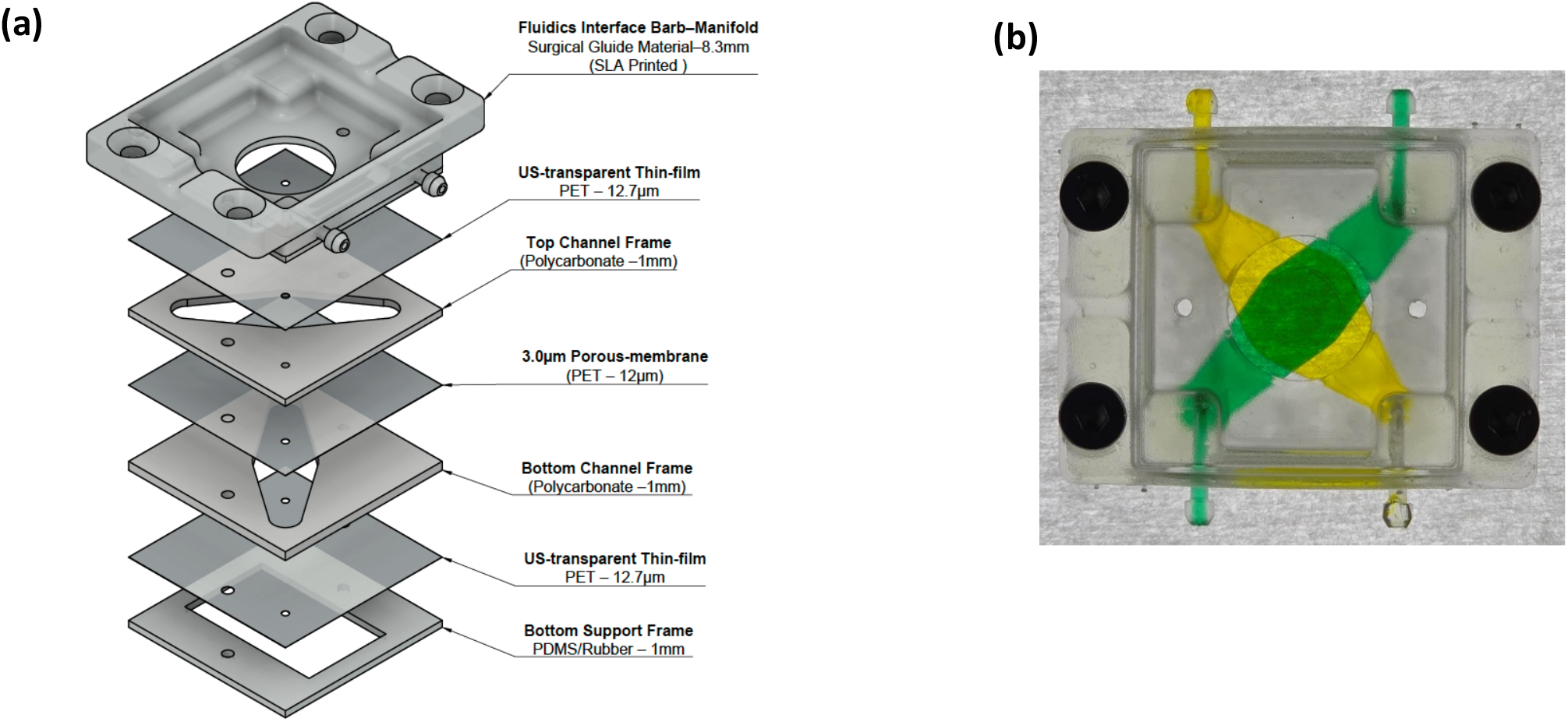
**a)** Structural architecture of the UST-OoC device with the UST membranes; **b)** Assembled UST-OoC device with a diagonal sandwich channel architecture with a central overlapping region.

### Acoustic Characterization and Multiphysics Simulation of the UST OoC device

The UST OoC device’s acoustic window includes a top and a bottom 12µm PET films and a center hydrophilic porous-PET membrane (3µm pore, 12µm thickness) with top and bottom channel heights of 1.28 mm (**Fig. 2b**). The |*t*| and |*r*| values of the UST OoC device were also measured similarly to the PET thin film measurement. **Fig. 3a** shows both the experimental data and COMSOL simulation of the |*t*| and |*r*| values for the complete three-membrane stack in an assembled device. Computationally, >99.9% transmission is present for all AOIs, with the highest |*t*| of 100% at normal AOI, and a minimum of 99.93% at 27.5°. Experimentally, the device maintains near-full transmission (>99.9%), with a small drop at 30° AOI to 99.23%, consistent with the theory. In any case, the net |*t*| remains at 99% or higher, significantly exceeding the current limitations of traditional cultureware, including available microfluidic OoC devices, which attenuate acoustic transmission by 60% or more at the first interface.^28^

**Figure 3:**
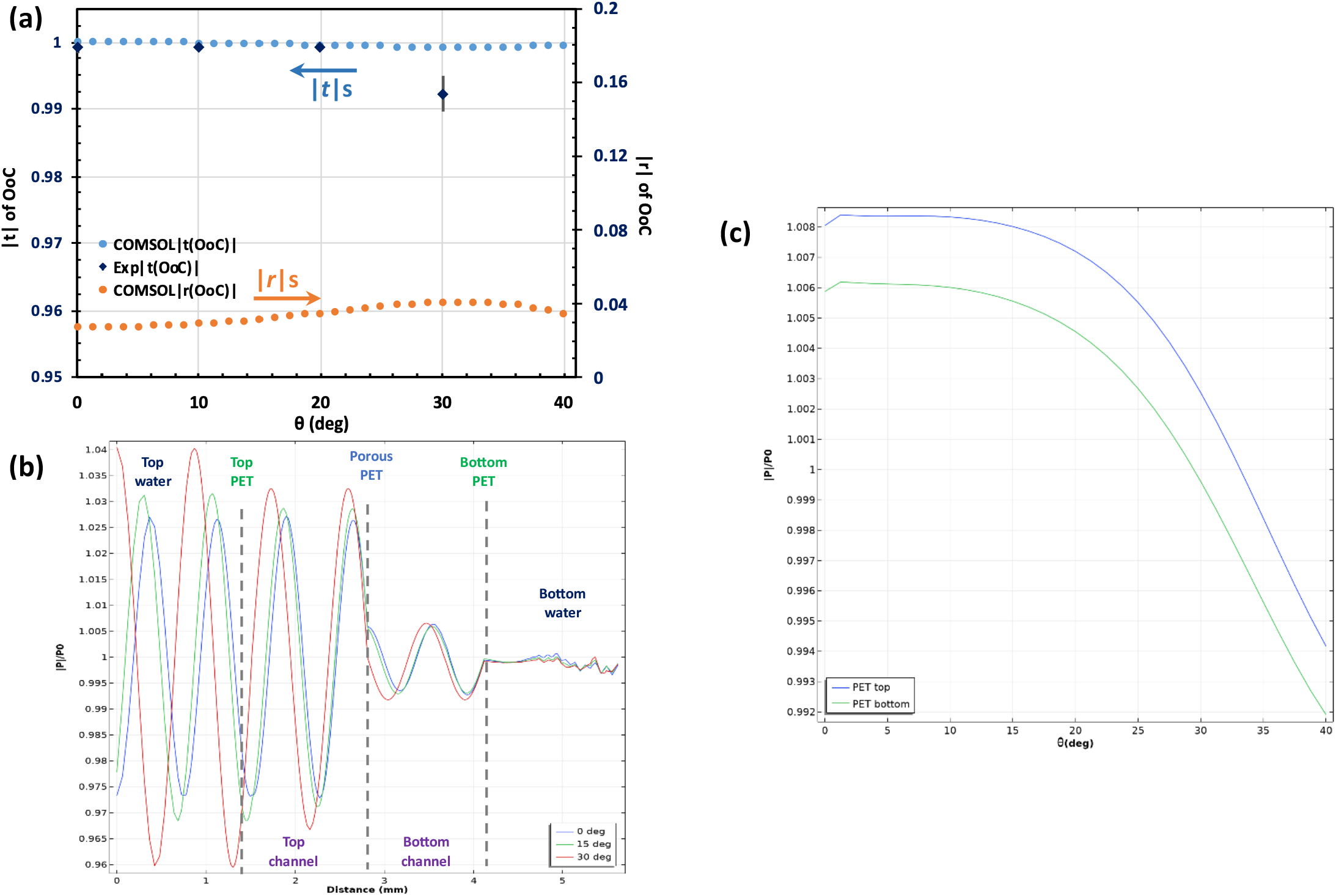
**a)** COMSOL-simulated |*t*| and |*r*| and experimental |*t*| for the OoC device stack at different incident angles, compared with experimental measurements; **b)** net acoustic field inside the OoC device for a plane wave incident at 0°, 15°, and 30°; **c)** normalized acoustic pressure inside the OoC device at 0°, 15°, and 30°, obtained by COMSOL simulation for a 1 MHz incident wave; **c)** normalized acoustic pressure at 2 µm above and below the PET membrane for 0°-40° angles of incidence. (P0 is the amplitude of the free-space US wave).

To better understand the acoustic field inside the OoC device, especially at the PET membrane surface where cells are cultured, we performed a Multiphysics COMSOL simulation, as measuring the acoustic field with a needle hydrophone inside the device is difficult and can itself perturb the acoustic field as a boundary. A simple plane US wave, instead of a focused beam, was used to gain qualitative understanding of the acoustic field (**see Fig. S2;** supplementary materials for details). The acoustic pressure amplitude I*P*I was normalized to the incident wave amplitude *P*_*O*_ along the height of the device and plotted for incident angles of 0°, 15°, and 30° (**Fig. 3b**). It was noted that standing waves can form inside the device due to the reflections of the bottom and middle PET membranes. The top channel shows greater energy variation due to positive interference between the two thin films. However, acoustic pressure variation is less than 3.5% due to the thin films’ low reflectivity, with a standing wave ratio (SWR) of 1.06, indicating accurate acoustic dose delivery inside the device. Furthermore, **Fig. 3c** shows that the acoustic field near the middle cell-culture PET membrane varies by less than 1% for AOI from 0 to 40 °, further confirming accurate acoustic dose delivery to the cells.

### Physical and Acoustic Characterization of submicron C_3_F_8_Gas-bubbles (sMBs)

Lipid-shelled gas bubbles were prepared using a Definity-like lipid blend and activated using amalgamation. We characterized the bubbles using Multi-angle Dynamic Light Scattering (MA-DLS). The resulting bubbles had a mean DLS hydrodynamic diameter of 806.1±92.5 nm (back-scatter) and a distribution-based polydispersity index of 0.0354±0.0062 AU (intensity-weighted percentage), with an average concentration of (2.045±0.560) × 10^10^ particles per mL and no further post-processing of the bubbles after activation (**Fig. 4a, b)**. The measured zeta potential in buffer was −19.67±1.05 mV.

**Figure 4:**
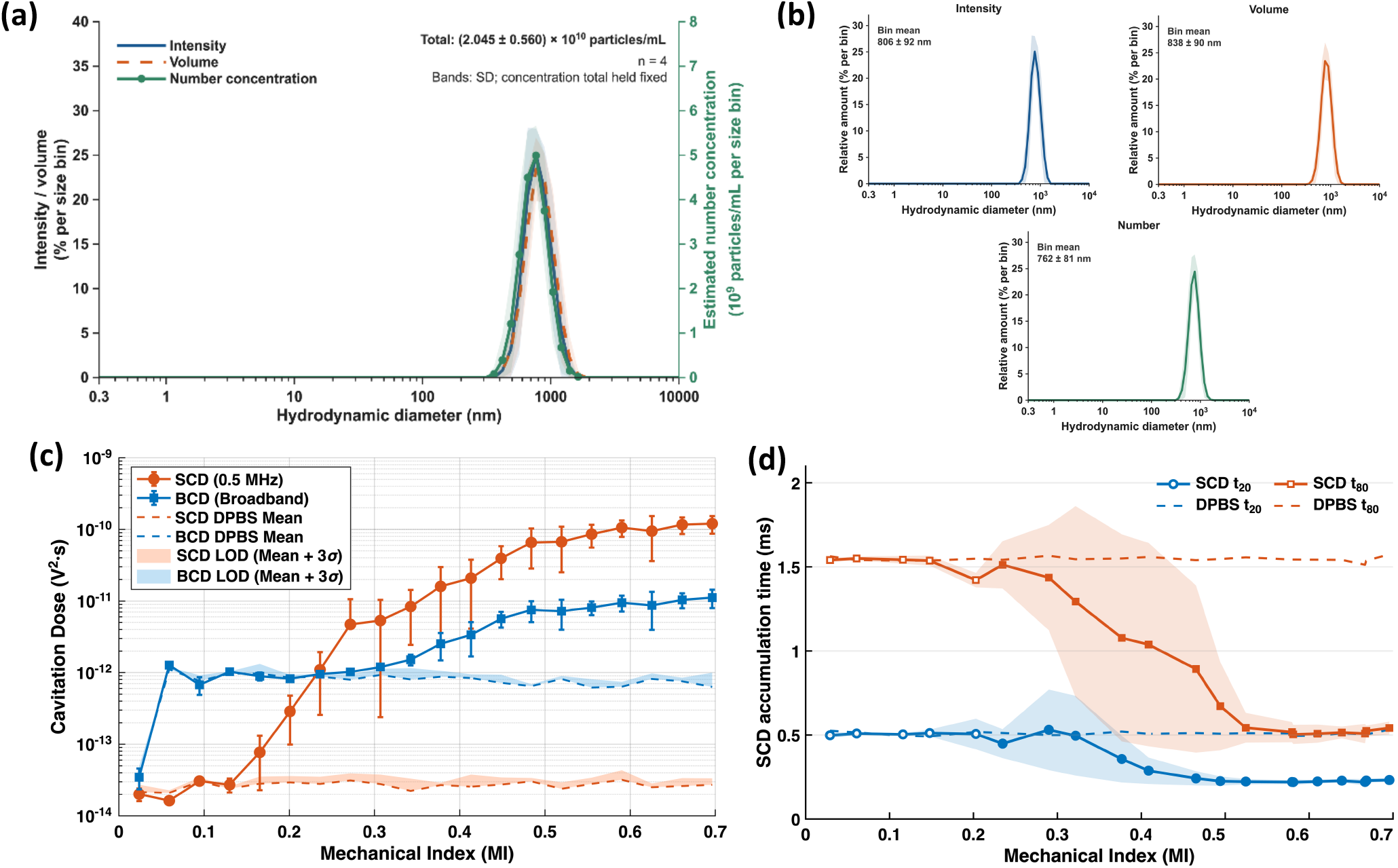
**a**. Intensity-weighted size distribution shows a submicron population measured by dynamic light scattering, and MADLS was used for concentration measurement; **b**. Size characterization of sub-micron microbubbles and their distribution based on Intensity, Volume, and Number; **c**. Acoustic dose escalation at a 1:4000 bubble dilution, measuring Subharmonic Cavitation Dose (SCD) and Broadband Cavitation Dose (BCD) across 0.029 to 0.696 MPa (MI 0.029-0.696 at 1 MHz). Plotted values are band-mean spectral densities, reported as mean ± SD across three replicates. Dashed curves show the degassed DPBS background; shaded regions span from the DPBS mean to mean + 3a, with a calculated over three groups of ten acquisitions. **d**. Times to reach 20% (T_20_) and 80% (T_80_) of retained subharmonic energy as a function of mechanical index. Shading shows ±SD; dashed curves display processed degassed DPBS. Time 0 ms marks the pulse start point.

For acoustic characterization, the sMBs were diluted at a rate of 1:4000, matching the dilution rate of the commercial Definity microbubble clinical dose. To evaluate the acoustic response to US, the sMBs were flown through the device’s apical channel at a fixed flow rate, underwent a mechanical index (MI) sweep across 20 ascending doses at 1 MHz (0.029 to 0.696), and passive acoustic emissions were collected and processed as previously described.^53^ **Fig. 4c** shows a subharmonic signal onset of 0.13 MI, and a broadband noise onset of ~ 0.3 MI. At MI 0.235, SCD was 9.8-fold above DPBS, whereas BCD remained near the reference level. Increasing MI from 0.235 to 0.696 increased SCD and BCD by approximately 418-fold and 13.6-fold, respectively. This rise was accompanied by earlier accumulation of subharmonic energy; the interval between T_20_ and T_80_ narrowed from 1.064 ± 0.064 to 0.310 ± 0.028 ms (**Fig 4d**). The central 60% of retained subharmonic energy therefore accumulated in an interval approximately 71% shorter, despite the unchanged 2-ms excitation.

Cumulative-energy measurements have previously been used to resolve differences in emission timing between lipid-bubble formulations^54^; Herein, we note that the same formulation exhibited a pronounced shift in timing across the pressure series. This dependence is especially relevant to bubble-cell interactions, where changes in pulse structure have been linked to differences in endothelial membrane poration and calcium signalling under flow.^39^ Our measurements show that the temporal distribution of bubble emissions also changes when pressure is varied at a fixed pulse duration. Further escalation produced little additional shift in T_80_. Across MI 0.525-0.696, mean T_80_ remained approximately 0.50−0.54 ms while SCD increased another 3.07-fold. Between these endpoints, subharmonic energy recorded before and after 0.5 ms both increased approximately threefold. Thus, the higher-pressure response grew through a broadly proportional increase in early and later emissions, with comparatively little change in their relative timing.

### Targeted Disruption of an Accelerated Caco-2 Barrier in the UST OoC using FUS^sMB^

FUS and gas bubble (FUS^GB^) is a promising approach that can transiently open the BBB (BBBO) at targeted brain locations for treating different neurological diseases.^9,55,56^ It has seen rapid adoption in the clinic, with 34+ trials across Phase I and II and multiple in Phase III.^57^ However, complications are still frequently reported, and a well-characterized precision in vitro model for the FUS^GB^ procedure remains unavailable, limiting the full understanding and optimization of the procedure, its regimens, and their safety profiles^58^.

To further assess the utility of the UST OoC device for therapeutic US studies in vitro, we performed sMB-assisted FUS (FUS^sMB^) disruption of an accelerated 3-day Caco-2 barrier as a proof of concept to demonstrate the targeted nature of the disruption within the UST OoC device platform. Caco-2 was chosen as a surrogate model because it exhibits strong paracellular barrier properties such as tight junction formation indicated by high-TEER (**Fig. 5a**) and expression of ZO-1 and occludin, as well as demonstrating key transport markers for glucose and efflux markers such as GLUT1 and P-glycoprotein (**Fig. 5b**).^59^ Among multiple mechanisms of FUS^GB^ BBB opening (e.g., tight junction disruption,^60^ efflux pump inhibition,^61^ caveolin-mediated transcytosis^62^, etc.), it is generally acknowledged that tight junction disruption to open the paracellular transport route is the main mechanism for reversible BBB opening, which makes Caco-2 a plausible surrogate model for studying FUS^sMB^ opening of the BBB *in vitro*, especially as an initial model for validation of the platform. The TEER value of the barrier in the Transwell insert reached ~ 1200Ω x cm^2^, which is higher than the 900Ω x cm^2^ TEER threshold needed for paracellular barrier studies of small molecules (< 500 Da) and larger IgG-class molecules (> 500 Da).^63^

**Figure 5:**
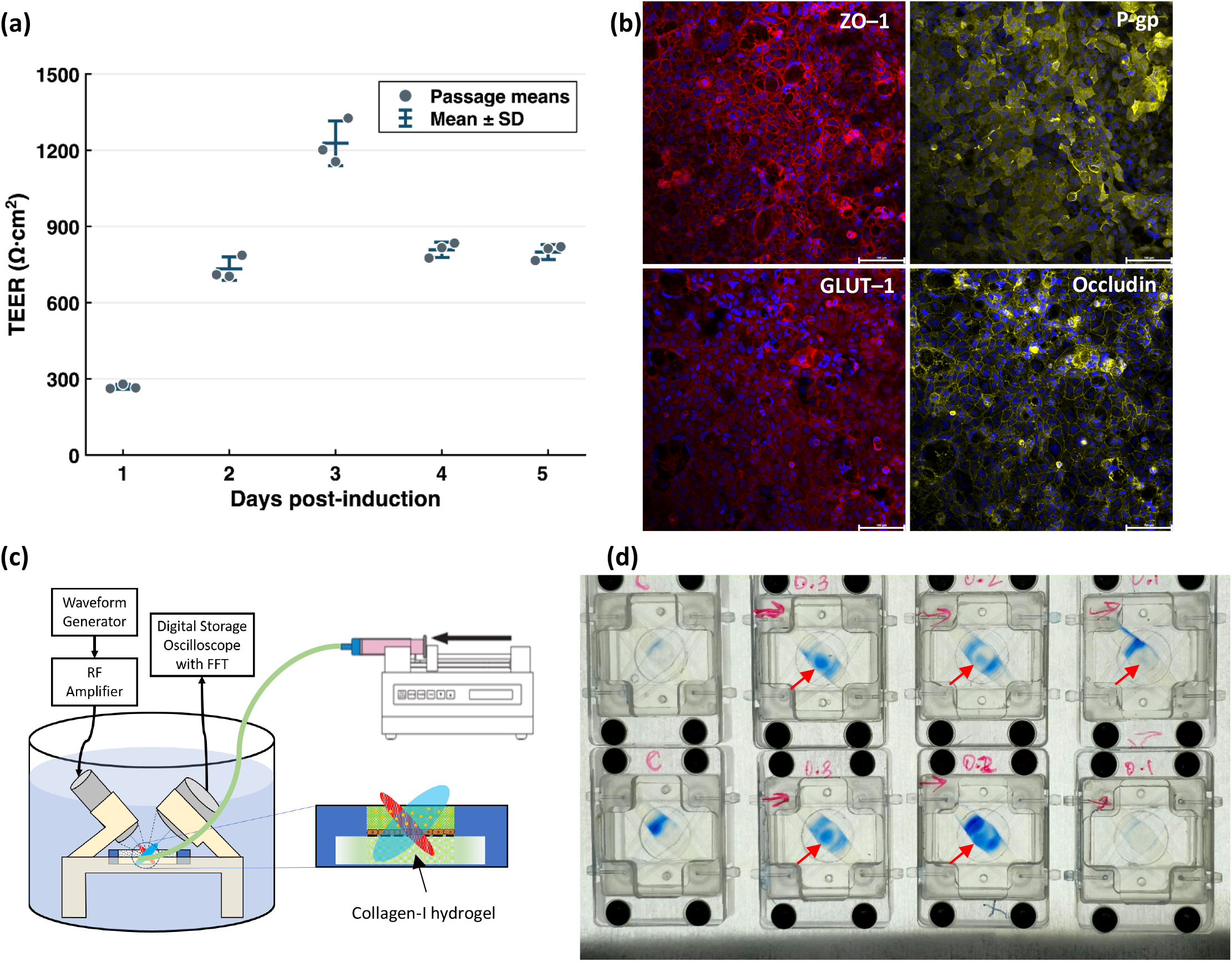
**a**. Transwell insert TEER values of the accelerated Caco-2 barrier over time; **b**. Immunofluorescence staining of tight junction proteins Occludin and ZO-1, as well as the GLUT1 transporter and P-gp efflux pump; **c**. schematic of the in vitro FUS BBBO setup. The bottom channel is filled with Collagen I hydrogel for the in vitro EBA assay; **d**. Preliminary results of acoustic dose (0, 0.1, 0.2, and 0.3 MPa) dependent targeted barrier opening by FUS. Red arrows indicate the targeted opening location. A leaky barrier at the edge of the channel was also observed.

An in vitro Evans Blue-Albumin (EBA) assay was developed to visualize targeted barrier disruption with spatial resolution. Evans Blue was conjugated to BSA hsV at a 3:1 molar ratio to form an EB–Albumin complex (~70 kDa) as the colorimetric marker for barrier opening. This ratio was chosen to minimize free EB dye in the tracer solution to <0.23%.^64^ FUS BBBO was performed at 1 MHz, PL 3 ms, PRF 1 Hz for 120 sec at 0, 0.1, 0.2, and 0.3 MPa while perfusing the top channel with a 1:4000-diluted sMB solution. After BBBO, the apical channel was incubated with the tracer complex and fixed. The Basolateral channel was loaded with 4 mg/mL Collagen I hydrogel to mimic the brain parenchyma, and the EBA was fixed to the Collagen I hydrogel after incubation. **Fig. 5d** shows the bright-field images of the disrupted barriers at different acoustic pressures. No targeted barrier disruption was observed in the control devices. One of the two devices showed targeted barrier disruption at 0.1 MPa. All barriers showed targeted disruptions at 0.2 and 0.3 MPa, with higher tracer intensity at 0.3 MPa. These results demonstrate, for the first time, acoustic dose-dependent targeted FUS barrier disruption *in vitro*.

In addition to targeted barrier disruption, the EBA assay also revealed random leaky barriers at the channel edges. The cause of the leaky barriers at the edges is still unclear and is speculated to arise from mechanical stress at the edges during device handling, further exacerbated by the adhesive type used to bond that interface, which has been documented to aggregate cells in clumps rather than allow monolayer formation. ^65^ It is under active investigation.

## Conclusion

In this study, we report the development of an UST OoC device for in vitro modeling of FUS-mediated BBBO for brain drug delivery. The acoustic transmission coefficients of a 12-µm-thick PET thin film at 1 MHz were theoretically calculated and experimentally characterized as 100% for AOIs up to 30°, with a slight 1.68% drop at 40°. An UST OoC device was fabricated using the PET thin film to form a top and a bottom UST window. COMSOL simulations were used to understand the acoustic field distribution within the device. The top channel showed higher but minimal acoustic pressure variation than the bottom channel (i.e., < 3.5%). An accelerated Caco-2 model was developed to reach a TEER of ~1200O x cm^2^ in 3 days for barrier disruption studies. The acoustic dose characteristics of the fabricated sMBs (~800 nm) at a 1:4000 dilution showed clear dose-dependent SCD and BCD signals, with onset thresholds of ~0.13 MI and ~0.3 MI, consistent with the in vivo therapeutic MI values used for FUS BBBO in the literature.^66^ Finally, we developed an in vitro EBA assay and observed acoustic dose-dependent targeted FUS barrier disruption. We are further optimizing the in vitro EBA assay conditions to generate additional data for statistical analysis and in vivo correlation. Overall, the results reported in the manuscript could serve as the first critical step toward quantitative in vitro modeling of FUS BBBO to understand the mechanisms of the procedure, as well as large parameter-space optimization, to develop novel procedure conditions for brain-drug delivery to treat different neurological diseases. We also envision direct or adapted broader applications of the UST OoC platform to study other therapeutic US applications, such as sonothrombolysis^67–69^, sonoporation/sonopermeation^70–72^, sonodynamic therapy^73^, US-triggered drug release^74,75^, etc.

## Experimental Methods

### Materials

Eagle’s Minimum Essential Medium (EMEM) (#30-2003) and Dimethyl sulfoxide (DMSO) (4-X) were purchased from ATCC. Fetal Bovine Serum heat inactivated (#A5669701), Nunc™ EasYFlask™ (156382), Penicillin-Streptomycin 10kU/mL (#15140122), HEPES (4-(2-hydroxyethyl)-1-piperazineethanesulfonic acid) (15630106), DPBS (10X), with calcium, magnesium (14-080-055), DPBS (1X), with calcium, magnesium (14040216), DPBS (10X), without calcium, magnesium (14200166), rMAb ZO-1 (MA5-46951), rMAb GLUT-1 (MA5-31960), Mab P-glycoprotein (MA5-13854), rMAb Occludin (740006M), Alexa Fluor Plus Donkey-Ms-488 (A32766TR), Alexa Fluor Plus Donkey-Rb-647 (A32795TR), FITC-Phallodin (A12379), Pierce™ 16% Formaldehyde (w/v), Methanol-free (PFA; 28906), DMSO, HPLC grade, 99.99+% (042780.AK), Live cell Imaging Solution (A59688DJ), Trypsin-EDTA 0.25% (25200-072) and Collagen I, rat tail (A1048301) were purchased from ThermoFisher Scientific (Waltham, MA).

Dulbecco’s Modified Eagle’s Medium (DMEM) 10-013-CV, Collagen I, Rat Tail low concentration (354236), Corning® Collagen I, High Concentration, Rat Tail (354249), Corning® MITO+ Serum Extender (355006), Corning® Bio-Coat® Poly-D-Lysine (354210), 100x Non-essential Amino acids (NEAA) (25-025-CI), and 100x Antibiotic-Antimycotic (30-004-CI) were kindly provided as a gift from Corning Inc. (Tewksbury, MA)

1N Sodium hydroxide (1N NaOH; S2770-100ML), Sodium butyrate (B5887-250MG), Evans Blue (E2129), Bovine Serum Albumin Fraction V, protease-free (3117332001), Triton™ X-100 (T8787), Propylene glycol (W294004-1KG-K), Sodium phosphate dibasic heptahydrate (S9390-100G), Sodium phosphate monobasic monohydrate (S9638-25G), Chloroform (C2432-1L), and Glycerol (G7893) were purchased from Millipore Sigma (St. Louis, MO).

1,2-dipalmitoyl-sn-glycero-3-phosphate (sodium salt) (DPPA, 16:0 PA; 830855P-25mg), 1,2-dihexadecanoyl-snglycero-3-phosphocholine (DPPC, 16:0 PC; 850355P-200mg), and 1,2-dipalmitoyl-sn-glycero-3-phosphoethanolamine-N-[carbonyl-methoxy(polyethylene glycol)-5000] (ammonium salt) (MPEG5000-DPPE, 880200P-200mg) were purchased from Avanti Polar Lipids (Alabaster, AL). Normal Goat Serum (005-000-121) was purchased from Jackson, IL. Vibrance® Antifade Mounting Medium with DAPI (H-1800-2) was purchased from Vector Labs (Newark, CA). Acetic acid 20 mM (5079) was purchased from Advanced Biomatrix. VWR® Spinbar® Micro Stir Bars Polygon 12mm x 3mm PTFE coated (58948-397) was purchased from VWR. Tygon SPT-3350 1/16” ID x 1/8” OD (EW-50113-90) tubing was purchased from Cole-Parmer (Vernon Hills, IL). Venor®GeM qOneStep (11-91025) used with Venor® Mycoplasma Extraction (56-3010), and Venor® Bacteria qPCR (12-2025) were purchased from Minerva Biolabs (Delaware, USA). NutriFreez® D10 Cryopreservation Medium (05-714-1B) and Microsart® ATMP Sterile Release Detection Kit (SMB95-1007) were purchased from Sartorius (Gottingen, Germany). Moisture-resistant polyester film (PET) (8567K102), all polycarbonate raw stock, end mills, taps, and bits were purchased through McMaster-Carr (Elmhurst, IL). Surgical Guide Resin (RS-F2-SGAM-01) was purchased through Formlabs (Somerville, MA). EVOM Manual (EVM-MT-03-02) and STX4 Electrodes (EVM-EL-03-03-01) were purchased from World Precision Instruments (Sarasota, FL). 2 mL vials, snap-top, Type I RSA Glass, 11mm snap ring (9509C-WCVRS), and Custom 11mm Crimp Aluminum cap, Septa were purchased from MICROSOLV (Leland, NC). Polyester (PET) Membrane Filters, Transparent, 3.0µm (1300032) were purchased from Steriltech (Auburn, WA). Pressuresensitive adhesives (PSA’s) ARseal™ MH94119 and ARcare® 90445Q were kindly provided as a gift from Adhesive Research (Glen Rock, PA)

### Cell Culture, Maintenance & Accelerated Differentiation

Caco-2 (HTB-37) (P-17, Lot# 70064029; ATCC) cells were purchased from the American Type Culture Collection and cultured in EMEM with 20% FBS, without antibiotics, and passaged routinely with 0.25% Trypsin-EDTA. Cells were then expanded in EMEM with 10% FBS with no antibiotics until P17-26 (P43) and cryopreserved as recommended by ATCC. Cells were then transitioned to DMEM + 10% FBS + 1X Anti-Anti at thaw, expanded for one passage, and frozen down in NutriFreez® D10. Cells were routinely surveilled for Mycoplasma and Bacterial contamination using qPCR-based testing. Before cryopreservation, cells were tested for mycoplasma, total bacterial, and fungal contamination using qPCR-based testing.

For all functional assays, cells were used at P17-29 (P46) and expanded up to 70-85% confluence in a T75 prior to use in DMEM + 10% FBS + 1X Anti-Anti.

A modification of a previously reported protocol from Corning for a 3-day barrier was used;^76,77^ For all experimental studies, cells were seeded in the device or inserts at a seeding density of 525,000 cells/cm^2^. Briefly, cells were dissociated from the flask, counted, and concentrated by spinning them down at 220*g* for 3 min; the cells were then resuspended and seeded in expansion medium DMEM + 1X NEAA + 10% FBS + 1X Anti-Anti + 20mM HEPES. Cells were then allowed to attach and expand. At 24 h, the medium was changed to differentiation medium (DM) consisting of DMEM + 1X NEAA + 1X Anti-Anti + 15mM HEPES + 1X MITO+ Serum Extender + 2 mM sodium butyrate. The cells were then cultured in the same medium with 24hr media changes until 96 h.

### UST-OoC Device Fabrication and Preparation

Polycarbonate (PC) sheet stock (1 mm thick) was machined using 2-axis milling (Roland MDX540) to produce the 26 mm x 26 mm apical and basolateral channel layers of the UST OoC device, including all features. All device membranes and adhesives were cut using low power and medium cut speed on a CO_2_ laser cutter (Versa Laser VL200, Scottsdale, AZ). The Fluidic Interface Barb-Manifold was printed in surgical-grade material at 50 µm layer thickness on a Form 3B+ SLA printer, then processed and washed in 99% isopropanol for 30 min, and cured on a Form Cure (Formlabs, Boston, MA). The PC layers were then cleaned with 70% ethanol, followed by 100% ethanol, and dried for at least 6 hours, up to overnight, in a 65°C dry oven. All assembly was performed in an ISO 6 cleanroom. Briefly, the device was bonded using the same top-down stack-up shown in **Fig. 2a**. The SLA printed Fluidic Interface Barb-Manifold was plasma-treated using oxygen plasma (50W, 25sccm O_2_ flow rate, 400 mTorr vacuum on Oxford RIE Plasma Lab 80Plus) for 1 min to improve surface adhesion and bonded with pre-cut PSA adhesive at all membrane-PC interfaces (MH94119) into the complete UST OoC device using an assembly jig consisting of a 1.5mm dowel pin with a PC base. The two UST PET thin films were pre-bonded to the PSA (90445Q) to improve membrane handling during assembly, and fluidic vias were biopsy-punched before the stack-up assembly. The completed device was then disinfected with 70% ethanol, followed by 100% ethanol, and incubated overnight (up to 24 h) at 50°C to drive out residual ethanol vapor.

The assembled and disinfected UST OoC device was then coated with Collagen I (rat tail) at 10 µg/cm^2^ in 20 mM acetic acid and incubated at RT in the BSC for 2 h, then washed twice with PBS, followed by seeding medium once and seeding cells.

### Immunofluorescent staining (IF) of Caco-2 and Imaging

Cells were seeded onto RSA chamber slides at 10 µg/cm^2^, matching the seeding density of the device/inserts, then fixed and stained using the standard protocol. Briefly, at 96 h of differentiation, cells were fixed in freshly prepared 4% PFA for 20 min, then rinsed 3X with DPBS^Mg+/Ca+^. They were then permeabilized with freshly prepared 0.25% Triton X-100 for 10 min and washed 5X with 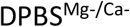. They were then blocked for 1.5 h in the same blocking buffer as the secondary Ab host at 5%, then incubated with Primary Ab diluted in 3% of the same blocking buffer overnight. The cells were then washed 5X with DPBSMg−/Ca− and incubated with Alexa Fluor Plus Secondary Ab at a 1:1000 dilution in the 3% blocking buffer for 1 h at RT before coverslipping (#1.5) with antifade sealant containing DAPI. Samples were allowed to seal for at least 24 h before imaging on a Leica Stellaris 8 confocal using a white-light laser with a 405nm diode laser.

### TEER measurement of Inserts and Analysis

Cells were seeded in collagen-I-coated inserts as previously described. Three separate passage replicates with a total of n=3 per passage were cultured and differentiated for a total of n=9 replicates per biological replicate. TEER was measured 24 h after induction of differentiation with DM and at 24 h intervals thereafter. TEER was measured using an EVOM Manual with an STX4 electrode in a 24-well insert (0.33 cm^2^) with basolateral chamber media volumes of 300 µL and 850 µL; each insert was measured at two positions (n=2 technical replicates). TEER is reported as (R_sample_ − R_blank_) × A, where R_blank_ is a coated insert with no cells, and A is the area of the insert. Measurements were made according to the manufacturer’s recommendations.

### In Vitro Evans Blue Assay

Prior to the FUS^NB^ procedure, the basolateral channel of the UST OoC device was loaded with a 4mg/mL final-concentration gel as a capture matrix for the Evans Blue-Albumin tracer complex. Evans Blue-Albumin tracer complex was prepared as a 3:1 molar ratio of Evans Blue to Bovine Serum albumin heat-shock fraction V (BSA-hsV). Evans Blue and BSA-hsV were prepared as absolute solutions at their maximum solubility in LCIS and then combined to form the conjugated tracer. Briefly, high-concentration Collagen I (Bornstein and Traub Type I, Rat tail, 10-11.5mg/mL) was neutralized by mixing with 5 parts of ddH_2_O and 1 part of 10x DPBS^Mg+/Ca+^, 0.2 parts of 1 M HEPES, 0.2 parts of 1N NaOH, and the remainder with Collagen I to bring it to a final concentration of 4mg/mL of Collagen in that order on ice, with a pH between 7.2-7.5. The neutralized Collagen mixture was then exchanged with the media in the basolateral channel, allowed to polymerize, and the barrier was allowed to recover at 37°C for 1.75 to 2 h.

### Transmission (|t|) and reflection (|r|) coefficient measurements of PET thin films

An ultrasound system comprising a waveform generator (Siglent SDG 1032X), an RF amplifier (NP961, NP Technologies), and a 1 MHz FUS transducer (A303S, Olympus, Evident Scientific) was used to generate 1 MHz FUS pulses with a pulse length (PL) of 2ms and a pulse repetition frequency (PRF) of 1 Hz. The transducer measures 12.7 mm and has a Point Target Focus (PTF) at 15.2 mm. The 6 dB beam width at the focal point was 1.94 mm, and the 6 dB axial beam depth was 15.0 mm. A 1-mm needle hydrophone (Precision Acoustics, UK) was mounted to a 3D-printed fixture connected to a homemade X-Y-Z positioning system to measure the transducer’s acoustic field. A digital storage oscilloscope (Siglent SDS1104X-E) measured the acoustic waveform from the hydrophone, and the PNP of the field was calculated using the hydrophone’s calibrated sensitivity at 1 MHz (138 mV/MPa) [**Fig. S1 A; Supplementary Material**].

For |*t*| measurements, the hydrophone was placed at the focal point along the transducer axis (see Fig. 1b). A PC frame was mounted in a holder attached to the hydrophone fixture, positioned 1 mm in front of the hydrophone, corresponding to the position of the cells within the device. This PC frame holder could be rotated to achieve different angles of incidence. The PC frame carried the PET membranes and the complete device stackup, and allowed exchange of replicates for measurements.

For |*t*| measurement, the hydrophone was oriented perpendicular to the transducer axis. The PC frame with the PET film was placed at a 45° angle to reflect the FUS pulse toward the needle hydrophone. The PET’s center was approximately 9 mm from the transducer surface. The needle hydrophone was then adjusted to the reflected focal point, determined by the time delay between pulse emission and sound-wave detection. The reflected wave amplitude by subtracting the background reflected wave amplitude of the PC frame without PET film from the reflected wave amplitude with the PDMS film.

For both |*t*| and |*r*| measurements, a 100 mVrms signal was applied at the waveform generator, producing an acoustic PNP of 0.3 MPa at the transducer’s focal point. The transducer, PET film, and hydrophone system were submerged in a custom water tank filled with degassed DI water. Each tank wall was lined with 10-mm-thick US absorbing material (AptFlex48, Precision Acoustics, UK). For each film, three samples were measured in duplicate by rotating the film by 180° to account for any direction-dependent variation in the film’s bulk properties, and these measurements were used to determine the mean and standard deviation.

### Acoustic Characterization and COMSOL simulation of UST OoC device

Theoretical |*t*| of the 12-µm-thick PET membrane in water at different incident angles was calculated using **Eq. (3)** with the density and sound speed values listed in **Table 1**. The experimental |*t*| of the PET membrane and the UST OoC was measured as for the PET thin film, except that a new rotating holder was made for the OoC device, with the transducer focal point positioned at the center of the PET membrane and the needle hydrophone placed along the transducer axis 1 mm behind the device. Duplicate measurements of four devices each were used to obtain the mean and standard deviation.

For COMSOL simulation of the acoustic field inside the device, we used a 2D Pressure Acoustic model to solve the frequency-domain acoustic field. The model geometry and mesh are shown in **Fig. S2a**. Briefly, the Acoustic module was used to simulate the acoustic field in the device from a uniform incident plane-wave at different AOIs using the properties in **Table 1**; a horizontal periodic boundary condition was used; a physics-controlled mesh was generated, consisting of a top and bottom perfectly matched layer (PML) at the outermost boundary domain, and the remainder of the geometry followed the device’s actual stack structure and inter-layer distances as per the actual CAD model.

### Submicron bubble (sMB) Formulation and Fabrication

A Definity-like lipid blend was used to fabricate the bubbles. Briefly, 16:0 PA, 16:0PC, and MPEG5000-DPPE were weighed in a vial at a weight ratio of 6.0:53.5:40.5 and dissolved in pre-warmed propylene glycol at 50°C for 5-10 min until fully dissolved to form the lipid blend. The lipid blend is then combined in a compounding vessel containing pre-warmed phosphate saline buffer containing glycerol and propylene glycol. The final formulation contained aqueous buffer, propylene glycol, and glycerol at an 8:1:1 volumetric ratio, with a total lipid concentration of 0.75 mg/mL and final salt concentrations of 4.87 mg/mL NaCl, 2.34 mg/mL NaH2PO4·H2O, and 2.16 mg/mL Na2HPO4·7H2O. The mixture was stirred during hydration for 20 min at 55-56°C, then ramped to 70-72°C and held for 5-7 min. After cooling to room temperature, the bubble-precursor-containing lipids were dispensed into glass vials, crimp-sealed, and stored at 4°C until activation. Before activation, the headspace was exchanged with octafluoropropane (C3F8) through repeated evacuation and gas-refilling cycles using a syringe-needle apparatus. The gas-exchanged vials were then immediately activated using a VialMix™ (Lantheus, Inc.), and an intermediate 1:100 dilution was prepared before diluting to a final 1:4000 dilution in Live-cell Imaging solution (LCIS).

### MADLS Bubble Characterization & Analysis

Bubbles were activated as described earlier and diluted to a 1:100 dilution for all DLS characterization. DLS size measurements were made in a 12 mm square polystyrene cuvette (DTS0012, Malvern Panalytical) using back-scatter measurement. Briefly, 1mL of the 1:100 bubble dilution in LCIS was loaded into the cuvette, loaded into the instrument, and measured per the manufacturer’s instructions at 25’C. Zeta potential measurements were made using a disposable folded capillary cell (DTS1070) per the manufacturer’s instructions.

Data analysis of the size was performed by exporting the Intensity-weighted hydrodynamic diameter distributions, software-derived volume- and number-weighted distributions, and plotting them. All exported size bins were retained. Distribution-based Polydispersity Index (PDI) was calculated using the intensityweighted distribution using 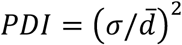, where σ is the standard deviation of particle diameters within the size distribution and 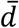 is the mean particle diameter for the distribution. Technical replicates were averaged within each vial and reported as the equally weighted mean ± standard deviation across the four vials (n=4) Particle concentrations were obtained using the same instrument at a 1:100 measurement dilution, and we reported the software-analyzed particle concentrations. The estimated number distribution of concentration per size bin was then computed by multiplying the average number fraction in each bin size by the measured average total concentration, as described before using *f*(*D*_*n*_) = *c f*(*D*_*n*_), where *c* was the mean stock particle concentration and *f*(*D*_*n*_) was the mean number fraction in diameter bin (n). All 70 native bins from the raw data. Shaded bands represent the standard deviation of number fractions multiplied by the fixed mean stock concentration.^78^ (**Fig. 4 a**,**b)**

### Acoustic Characterization in the UST OoC device & Analysis

Bubbles were activated as described, diluted in DPBS to 1:4000, and perfused through the UST OoC device at 300 µL/min in the apical channel; the basolateral channel was filled with LCIS and capped. A pressurecompensation vessel containing the same DPBS used to dilute the bubbles was placed at the end of the exit tubing to ensure consistent hydrostatic pressure at the channel level. The channel was positioned at the overlapping focal regions of a single radial-element 1 MHz transmitting transducer (Olympus A303S) and a single radial-element 0.5 MHz passive cavitation detector (Olympus A301S). A Python script using the VISA package (National Instruments) controlled the waveform generator and the oscilloscope from a bash terminal via SCPI commands, adjusting power levels and capturing raw waveforms from the oscilloscope. A 1 MHz, 0.2% dutycycle pulse was used at a pulse-repetition frequency (PRF) of 1 Hz.

Pressure was increased from 0.029 to 0.696 MPa across 20 points; at each point, 30 raw waveform acquisitions were taken per replicate, at a sampling depth of 70kpts with the trigger matched to the waveform generator (n=4). Degassed DPBS was measured over the same pressure series.

Signals were processed in MATLAB as previously described^79^, and a Tukey window with a taper r=0.1 was applied. FFT magnitudes were averaged across captures, divided by the processed waveforms, squared, and divided by the frequency-bin spacing^80^. Band-mean spectral densities covered 0.45-0.55 MHz for subharmonic cavitation dose (SCD) and 0-2.5 MHz for broadband emissions (BCD), excluding ±0.05 MHz around 0.5, 1, 1.5, and 2 MHz. Results are mean ± SD from three vials. DPBS reference curves and mean + 3 SD boundaries (shaded regions; **Fig. 4 c**) were calculated from n=3 measurements. For temporal analysis, the subharmonic band and its negative-frequency counterpart were retained, and inverse Fourier transformation was used to recover the bandlimited voltage waveform. The squared voltage was averaged over the 30 captures before cumulative summation and normalization. T_20_ and T_80_ were defined as the first times at which 20% and 80% of the retained energy were reached, respectively, following the method adapted from Pouliopoulos et al.^54^ Timing was referenced to the common −1000 µs acquisition coordinate, preserving the same processing as the dose-escalation. Results were reported as mean ± SD across three vials, with DPBS processed and displayed separately (**Fig 4d**).

### FUS^sMB^ Disruption in the UST OoC-device

Cells were cultured in the device, and Collagen was gelled in the basolateral channel as described earlier. A 1:4000 dilution of bubbles in LCIS was flowed into the device’s apical channel, and disruption was performed using a 1 MHz, 3ms pulse at a pulse-repetition frequency (PRF) of 1 Hz; a total of 120 pulses were delivered at 0.3MPa. After the FUS^sMB^ procedure, we loaded the Evans blue-albumin tracer into the apical channel and incubated it for 30 min at RT before washing with DPBS^Mg+/Ca+^ and fixing with 4% PFA, then imaging on an illuminated background.

## Data availability Statement & Materials availability

All data are included within the article and/or the supplementary information with this manuscript. The devices generated in this study will be available on request, but we may require payment and/or execution of a materials transfer agreement (MTA).

## Acknowledgements

This study was supported by the UA SensorLab Seed Grant (J.G. and F.Z.), UA BMS CRP grant (J.G. and R.J.G.), UA ORP Bridge Funding (J.G. and R.J.G.), and the Center for Applied NanoBioscience and Medicine.

The authors thank the Helios Education Foundation and the Translational Genomics Institute for supporting undergraduate and graduate students and programs. The authors also thank all members of the Center for Applied NanoBioscience and Medicine for their support and contributions. We extend our gratitude to Drs. Shenfeng Qiu and Kurt Gustin, director of the Biomedical Imaging Core (BIC) at the UA College of Medicine-Phoenix, for providing confocal imaging services, to Dr. Timothy Marlowe, director of the Molecular Discovery Core at the UA College of Medicine - Phoenix, for providing access to fluorescence plate reader services and Dr. Zhicheng Deng at the Phoenix Children’s Hospital Research Center for providing access to the DLS instrument.

## Author Contributions

J.G., F.Z., and N.M.F. contributed to conceptualization of the study; N.M.F., R.J.G., F.Z., and J.G. contributed to data curation and study design; N.M.F., N.H., L.K., S.A., B.T., J.G. contributed to methodology; N.M.F., N.H., L.K., B.L., S.A., B.T., and M.W.B. contributed to investigation and data acquisition. N.M.F., B.L., and J.G. contributed to visualization. N.M.F. and J.G. contributed to formal analysis and writing - original draft of this manuscript. N.M.F., N.H., L.K., B.L., S.A., B.T., M.W.B., R.J.G., F.Z., and J.G. contributed to writing - review and editing of this manuscript. N.M.F. and J.G. contributed to supervision. J.G., R.J.G., and F.Z. contributed to project administration. J.G., R.J.G., and F.Z. contributed to funding acquisition.

## Notes

### Competing Interest Statement

J.G., F.Z., and N.M.F. are inventors on US Patent Application No.19/631,845, and PCT Application No. PCT/US24/49098. All other authors declare no competing interests.

