## Supplemental Materials for "Towards Quantitative *In Vitro* Modelling of Focused Ultrasound-mediated Blood-Brain Barrier Opening using an Ultrasound-transparent Organ-on-Chip Device"

**Jian Gu, Ph.D.**

### Towards Quantitative *In Vitro* Modelling of Focused Ultrasound-mediated Blood-Brain Barrier Opening using an Ultrasound-transparent Organ-on-Chip Device

**Nabhan M. Fakrudin, Nathan Han, Laith Kabbani, Brandon Lan, Suzan Aldekr, Baiju Thomas, Matthew W. Barrett, Rayna Gonzales, Frederic Zenhausern, and Jian Gu***

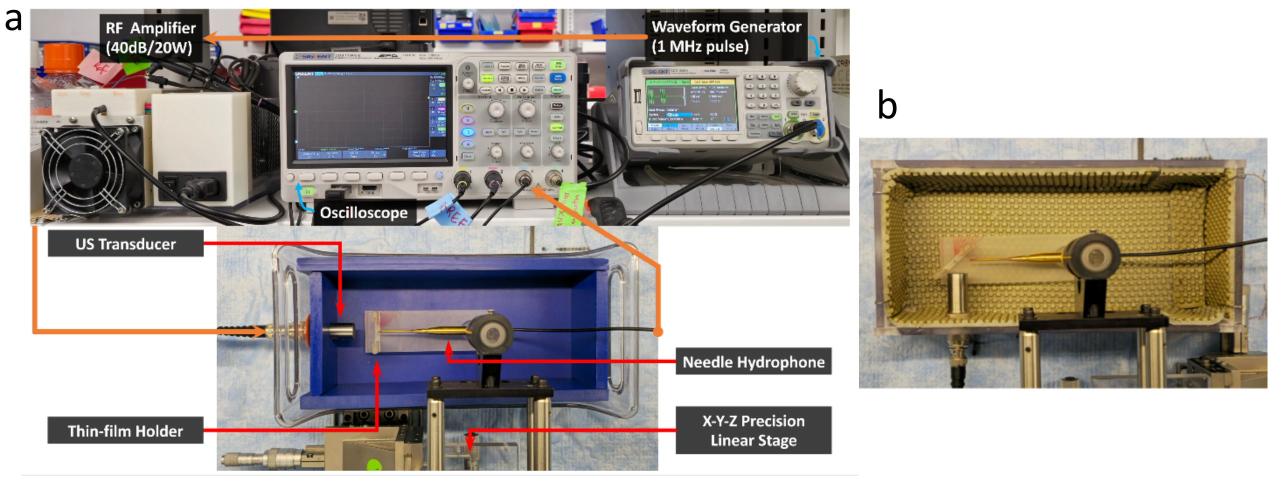


**Fig. S1.** An overview of the transparency characterization system and the fabricated tanks. (a) The US transparency characterization setup before measurement for transparency characterization. The thin-film holder is mounted with a screw at the pivot point to allow changes in the Angle of Incidence (AoI) of the US beam relative to the thin film. (b) Tank fabricated for reflection measurements, with AoI fixed at 45° for all reflection measurements due to limitations of the experimental setup.

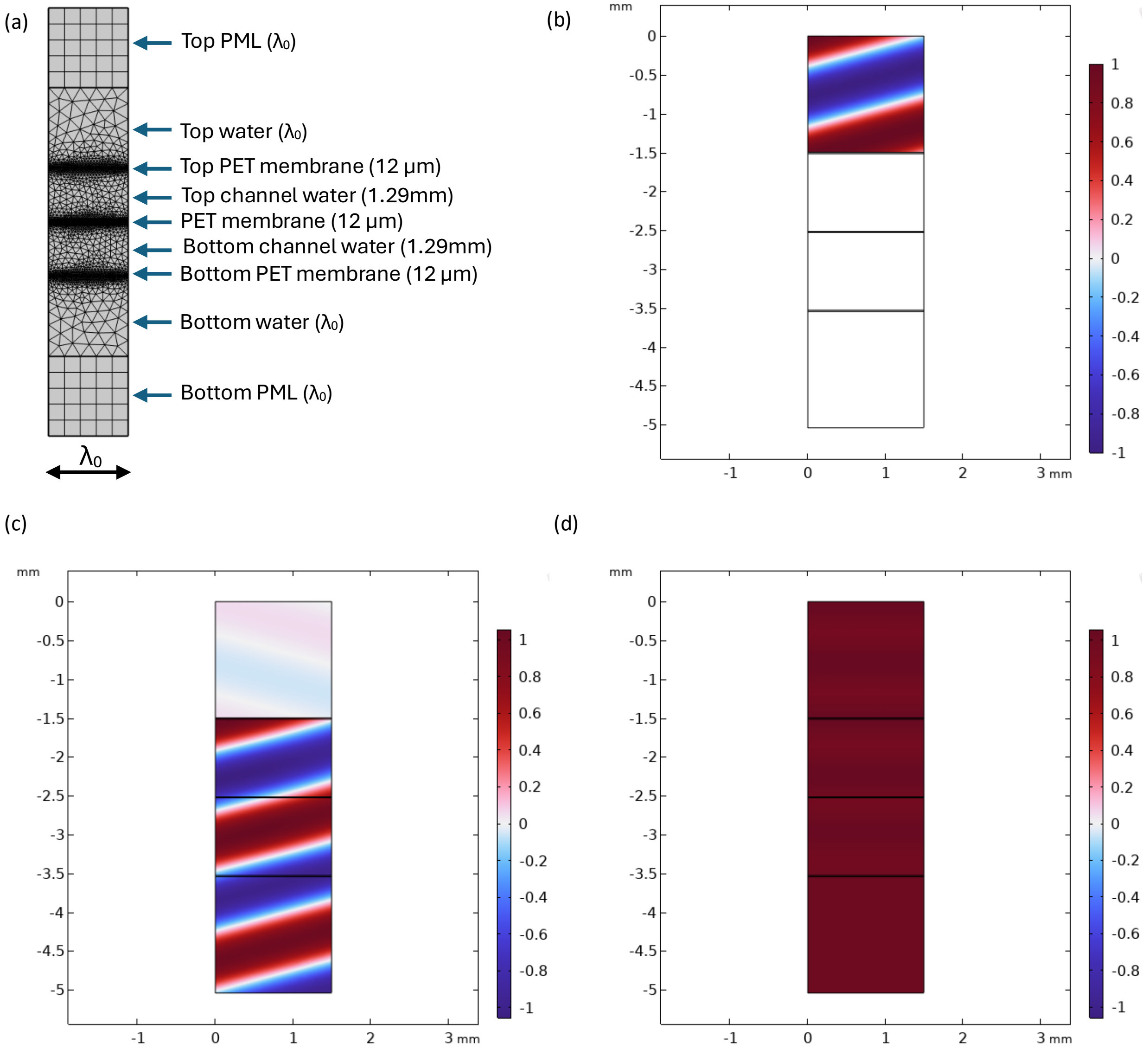


**Fig. S2** (a) mesh of the 2D acoustic pressure/frequency domain model; the width of the structure is the US wavelength in water ($\lambda_0$=1.497 mm), the heights are noted in parentheses; (b-d) normalized incident, scattered, and total acoustic field at a 15° incident angle.
